# Selection of start codon during mRNA scanning in eukaryotic translation initiation

**DOI:** 10.1101/2020.11.06.371484

**Authors:** Ipsita Basu, Biswajit Gorai, Thyageshwar Chandran, Prabal K. Maiti, Tanweer Hussain

**Affiliations:** Center for Condensed Matter Theory, Department of Physics, Indian Institute of Science, Bangalore-560012, India; Department of Molecular Reproduction, Development and Genetics, Division of Biological Sciences, Indian Institute of Science, Bangalore-560012, India

**Author notes:** These authors contributed equally to this work (Biswajit Gorai, Thyageshwar Chandran). Department of Biotechnology, National Institute of Technology-Warangal, Telangana-506004, India (Thyageshwar Chandran).

**Keywords:** ribosome, translation initiation, scanning, start codon, binding energy, molecular dynamics

## Abstract

During translational initiation in eukaryotes, the small ribosomal subunit forms a 48S preinitiation complex (PIC) with initiation factors. The 48S PIC binds to the 5’ end of mRNA and inspects long untranslated region (UTR) for the presence of the start codon (AUG). Accurate and high speed of scanning 5’ UTR and subsequent selection of the correct start codon are crucial for protein synthesis. However, the conformational state of 48S PIC required for inspecting every codon is not clearly understood. Whether the scanning or open conformation of 48S PIC can accurately select the cognate start codon over near/non-cognate codons, or this discrimination is carried out only in the scanning-arrested or closed conformation of 48S PIC. Here, using atomistic molecular dynamics (MD) simulations and free energy calculations, we show that the scanning conformation of 48S PIC can reject all but 4 of the 63 non-AUG codons. Among nine near-cognate codons with a single mismatch, only codons with a first position mismatch (GUG, CUG and UUG) or a pyrimidine mismatch at the second position (ACG) are not discriminated by scanning state of 48S PIC. In contrast, any mismatch in the third position is rejected. Simulations runs in absence of one or more eukaryotic initiation factors (eIF1, eIF1+eIF1A, eIF2ɑ or eIF2β) from the system show critical role of eIF1 and eIF2ɑ in start codon selection. The structural analysis indicates that tRNAi dynamics at the widened P site of 48S open state drives codon selection. Further, a stable codon: anticodon interaction prepares the PIC to transit to the closed state. Overall, we provide insights into the selection of start codon during scanning and how the open conformation of 48S PIC can scan long 5’ UTRs with accuracy and high speed without the requirement of sampling the closed state for every codon.

## 1. Introduction

Eukaryotic translational initiation is a complicated yet well-coordinated process. The preinitiation complex (PIC; the complex of 40S ribosomal subunit and initiation factors) binds to the 5’ end of mRNA and scans along until a start codon is encountered in the ribosomal P site [1,2]. In brief, the overall event starts with the initiation factors eIF1, eIF1A and eIF3 binding to the 40S subunit, which in turn facilitates the binding of charged initiator tRNA (Met-tRNAi) as a ternary complex (TC) with eIF2-GTP [3]. The 43S PIC thus formed is recruited to capped 5’ end of mRNA with the help of eIF4 factors [4,5,6], leading to the formation of 48S PIC.

Unlike in prokaryotes, eukaryotic genes have long 5’ UTRs that can exceed 1000 nucleotides; for instance, in humans, the maximum length reported is 2803 [7,8,9]. The median length of 5’ UTR is ~53 nucleotides in budding yeast and in the range of 53-218 nucleotides for higher organisms [9]. A recent finding shows that only one ribosome scans a 5’ UTR at a time in most human cells, and the length of 5’ UTR affects translation efficiency [10]. Hence, it is imperative to address how the ribosomal PIC can scan a long 5’ untranslated region with high speed and accuracy.

To recognize the start codon, the 48S PIC has to accurately inspect successive triplets of mRNA nucleotides entering the P site for complementarity to the anticodon of tRNAi at high speed. The process is unidirectional and in base-per-base mode [11]. The net speed of scanning was about 8-10 nucleotides per second in cell-free extract, and the scanning rate is expected to be even higher within the cell [11,12]. However, there is no clear understanding of the scanning mechanism and how the 48S PIC can scan at such a high speed with accuracy.

It is assumed that the 48S PIC would continue to scan the 5’ UTR in an open or scanning or (P_OUT_) conformation until it reaches the start codon. Recognition of the start codon leads to scanning arrest and conformational rearrangement to the closed state (P_IN_) of the 48S PIC [3,13]. Further, the downstream events in translation initiation are triggered following the release of eIF1, which is essential for the fidelity of start codon selection [1,2,3,14,15]. Alternatively, it is suggested that the 48S PIC may shuttle spontaneously between open and closed conformations during scanning [16,17,18]. Based on the latter model, the energetics of initiator tRNA binding to different near-cognate codons in the yeast 48S PIC in closed conformation was studied using atomistic molecular dynamics (MD) simulation and free energy calculation in the presence or absence of eIF1 and eIF1A to understand scanning [16,17]. The results indicated that eIF1 was primarily involved in discrimination against mismatches in the first and second positions of the codon, whereas eIF1A played a role in discrimination against near-cognate codons with third position mismatches in the P_IN_ state [16]. However, these results consider the energetics of codon discrimination only in a scanning-arrested P_IN_ state, which cannot be extrapolated to the open, scanning conformation of the 48S PIC because of the conformational differences in the two states (discussed below). Furthermore, the authors considered the coordinates of a closed-state 48S PIC without the tRNA_i_ as a model for the scanning state [16]; accordingly, it was assumed that no interaction between codon and anticodon exits in the open state. However, with the availability of a cryo-electron microscopy (cryo-EM) structure of a 48S PIC in its open conformation (py48S-open complex) [19,20], it became clear that the open conformation is significantly different from the closed one (py48S-closed complex). Importantly, codon-anticodon interaction does indeed occur in the open state of 48S PIC.

In the open conformation of 48S PIC, the 40S head moves upwards with respect to the body with attendant relaxation of rRNA helix 28, which connects the head to the body of the 40S. As a result, both the mRNA channel and the P site are widened, and the mRNA latch is opened.

The tRNA_i_ is positioned in the P site ~7 Å away from 40S body compared to that found in the py48S-closed complex. eIF1 is observed at its primary binding position at the P site in the open state. It undergoes subtle repositioning and deformation on the transition to the closed state to accommodate tRNA_i_ in the P_IN_ conformation [19]. Strikingly, the N-terminal tail (NTT) of eIF1A, which was observed in proximity to the codon: anticodon duplex in py48S-closed complex, could not be observed in the py48S-open complex, highlighting its role in stabilizing specifically the closed conformation of the 48S. Further, eIF2β contacts eIF1 in the py48S-open complex but moves away from eIF1 in the py48S-closed complex, indicating its role in stabilizing the open conformation. Given these striking differences in the conformation of 40S, eIFs and tRNA_i_ between open and closed states of the 48S PIC, the coordinates of the closed state of the 48S PIC without the tRNA_i_ do not accurately reflect the correct open conformation.

Moreover, the observation of codon: anticodon interaction in the structure of the py48S-open complex with an AUC start codon in the P site [19] indicated that the tRNA_i_ could inspect the incoming codon in the P site even in the open conformation. This led us to consider whether the 48S PIC in an open conformation can accurately recognize the start codon and discriminate against non-cognate codons while scanning the 5’ UTR. If this is indeed true, then the 48S PIC would not have to undergo a large conformational change (from open to closed state and back) to inspect every incoming nucleotide triplet in the P site explaining the high speed of scanning.

Therefore, in this study, we decided to regard the binding energy of the tRNA_i_ with each non-cognate codon relative to the cognate AUG start codon in the open conformation of the 48S PIC as a determinant of its frequency of selection as a start codon during scanning. A similar approach was used earlier to examine the selection of non-cognate start codons in the closed state [16]. Here, we report that the open conformation can recognize the start codon AUG and discriminate against most non-cognate codons. Recognition of AUG in the open state seems to prepare the 48S PIC to change its conformation to the closed state. Our studies also indicate that eIF1 plays a crucial role in codon selectivity in the open conformation of the 48S PIC. However, recognition of AUG as a start codon is still inaccurate owing to the failure to discriminate against codons (GUG, CUG, UUG) with a first base-pair codon: anticodon mismatch. Hence, the open conformation of the 48S PIC serves as an initial checkpoint for selection of start codon at which almost all non-cognate codons are rejected. A few near-cognate codons, which are accepted in the open state, can then be re-examined in a more stringent, second checkpoint, i.e., the closed conformation of the 48S PIC. Thus, our study provides novel insights into how the 48S maintains accuracy at a high rate of scanning by utilising the open state as a coarse selectivity checkpoint to reject all but a few of the possible codon: anticodon mismatches.

## 2. MATERIALS AND METHODS

### 2.1. Overall design of systems for molecular simulation dynamics

We based our analysis on the py48S-open-eIF3 complex (PDB ID: 6GSM) determined at 5.2 Å resolution [20]. Overall this structure is similar to the py48S-open complex reported previously (PDB ID: 3JAQ) [19] and has density for three nucleotides (AAU) of mRNA corresponding to ‘A’ at the −1 position and ‘AU’ in the +1 and +2 positions of the AUC codon in the P site, and weak density for mRNA observed throughout the mRNA channel. Hence, we decided to model the remaining nucleotides of mRNA in the mRNA channel for our calculations, as this would mimic the state when PIC is scanning the 5’ UTR. Since the mRNAs in the py48S-open-eIF3 complex (PDB-id: 6GSM) and py48S-eIF5N complex (PDB ID: 6FYY) [21] differ only in a single base (i.e., AUC vs. AUG codon at the P site), we have modeled the mRNA in the remaining mRNA channel based on the mRNA observed in the py48S-eIF5N complex (PDB ID: 6FYY).

It is computationally expensive to simulate the whole py48S-open-eIF3 complex, so we generated a simulation sphere of 40 Å radius centered on the center of mass (COM) of the ‘A35’ nucleotide of the anticodon (5’-CAU-3’) of tRNA_i_ at the P site. Previously, a 25 Å simulation sphere from the py48S complex was used for free energy calculations of the 48S PIC in the PIN state [16]. Since the POUT state has a widened mRNA channel, we decided to increase the radius of the simulation sphere to account for this feature, as well as to include more contributions from ribosomal components, tRNA_i_ and bound eIFs, namely eIF1, eIF1A, eIF2α and eIF2β.

### 2.2. Generation of near-cognate and non-cognate codons at the P site

To compare the relative binding energy of the tRNAi with different non-cognate codons in the P site with respect to AUG, 3-, 2- and 1-point mutations were made in the start codon. Thus, the only change in the simulation sphere is in the codon at the P site, while the rest of the atomic coordinates remain unchanged. The individual mutations of the start codon at the P site were introduced using the module “mutate_bases” of X3DNA software [22]. In order to study the effect of the eIFs, atomic coordinates of each factor individually, or combinations of eIFs, were excluded from the simulation sphere before the respective production run.

### 2.3. Molecular dynamics simulation protocol

Molecular dynamics (MD) simulations were carried out using the PMEMD module of the AMBER14 package [23]. The TIP3P water model [24] was used to solvate such that the solvation shell extends at least 15 Å in all directions from the solute. A requisite number of Na^+^ ions were added to the systems to maintain overall charge neutrality. The resulting structure contains ~188200 atoms making it one of the largest system simulated according to our knowledge. The Xleap module of the AMBER14 package was used to solvate and add the ions. AMBER ff14SB force field [25] was used to describe the interactions involving proteins, RNA and water. Joung-Cheatham ion parameters [26] was used to describe interactions involving ions. Energy minimization was then performed on the solvated systems for 3000 steps using the steepest descent method, followed by 3000 steps of the conjugate gradient method. The atomic coordinates in the simulation sphere was kept fixed to its initial structure, using a harmonic restraint of 500 kcal/mol/Å^2^ during this initial minimization. This minimization was followed by another conjugate gradient minimization by slowly reducing the force constant of the harmonic restraint on the ribosome from 20 to 0 kcal/mol/Å^2^ in five consecutive steps. The system was then gradually heated to a temperature of 300 K in two steps: first, the NVT ensemble was involved in heating from 0 to 100 K in 8 ps and then systems were heated to 300 K using the NPT ensemble in 80 ps. The solute particles were restrained to their initial positions using harmonic restraints with a force constant of 20 kcal/mol/Å^2^ during the whole heating process followed by 500 ps of equilibration run in NPT ensemble using a 2 fs time step for integration. This step was followed by 60 ns of NPT production run where the atoms in the residues of ribosome which are further beyond 30 Å from the center of the ribosome sphere will be harmonically restrained with 10 kcal/mol/Å^2^ force constant. The pressure was kept constant at 1 atm using the Berendsen weak coupling method [27].

The short-ranged vdW and electrostatic interactions were truncated within a real space cut off of 10 Å and the particle mesh Ewald (PME) method was used to calculate long-range electrostatic interactions. All bond lengths involving hydrogen atoms were constrained using the SHAKE algorithm. The simulation runs for a codon have been repeated four to six times with different initial velocities. The snapshots are generated using VMD [28].

### 2.4. MM-PBSA Free energy calculation

The MM-PBSA (MM: Molecular Mechanics; PB: Poisson Boltzmann; SA: Surface area) method was used to calculate relative free energies of binding using the MMPBSA.py [29] module of AmberTools14. The binding free energy is computed as ΔG_bind_ = ΔE_bind_ − TΔS_bind_, where ΔE_bind_ is the sum of the changes in the electrostatic energy and TΔS is the solute entropy term which has been neglected for the present calculation. ΔE_bind_ is expressed as ΔE_bind_ = ΔE_ele_ + ΔE_vdw_ + ΔE_int_ + ΔE_sol_, where ΔE_ele_ is the changes in electrostatic energy, ΔE_vdw_ is the non-bond van der Waals energy, ΔE_int_ is the internal energy from bonds, angles and torsions, and the contribution from the solvent is ΔE_sol_. ΔE_sol_ is the sum of the electrostatic energy, ΔE_es_ and the non-electrostatic energy, ΔE_nes_. ΔE_es_ is calculated using the Poisson–Boltzmann (PB) method and ΔE_nes_ is expressed as ΔE_nes_ *= γ*SASA + *β*, where *γ* = 0.00542 kcal Å^−2^ is the surface tension, *β* = 0.92 kcal mol^−1^, and SASA is the solvent-accessible surface area of the molecule. The time series of the binding free energy of the codon-anticodon complex was determined using gas-phase energies (MM) and solvation free energies following the Poisson Boltzmann model (PB/SA) analysis from snapshots obtained from the last 40 ns of total 60 ns of simulation trajectory and averaged over all the independent runs for each system.

## 3. RESULTS AND DISCUSSION

### 3.1 Relative binding energies of non- and near-cognate codons: A possible cue for codon selection

The scanning of the 5’ UTR by the 48S PIC primarily involves the anticodon of tRNA_i_ interacting with mRNA at the P site, actively encountering codon triplets probably with one, two, or three mismatches compared to the cognate start codon, AUG [1]. In order to understand the mechanism of codon selection in an open conformation of the 48S PIC, we carried out an array of comprehensive MD simulations of the core area of the py48S-open-eIF3 PIC (Fig. 1 and Supplementary Fig. S1). The simulations were carried out with cognate (AUG) and multiple non-AUG codons with one, two, or three mismatches (Fig. 2 and Supplementary Fig. S2). The relative free energy perturbations from the respective simulations provided insight into codon selection by the PIC in its open state. Simulations were also carried out by excluding one or more initiation factors from the system (eIF1, eIF1+eIF1A, eIF2β, or eIF2ɑ) to evaluate their roles in the mechanism of start codon recognition.

**Fig. 1:**
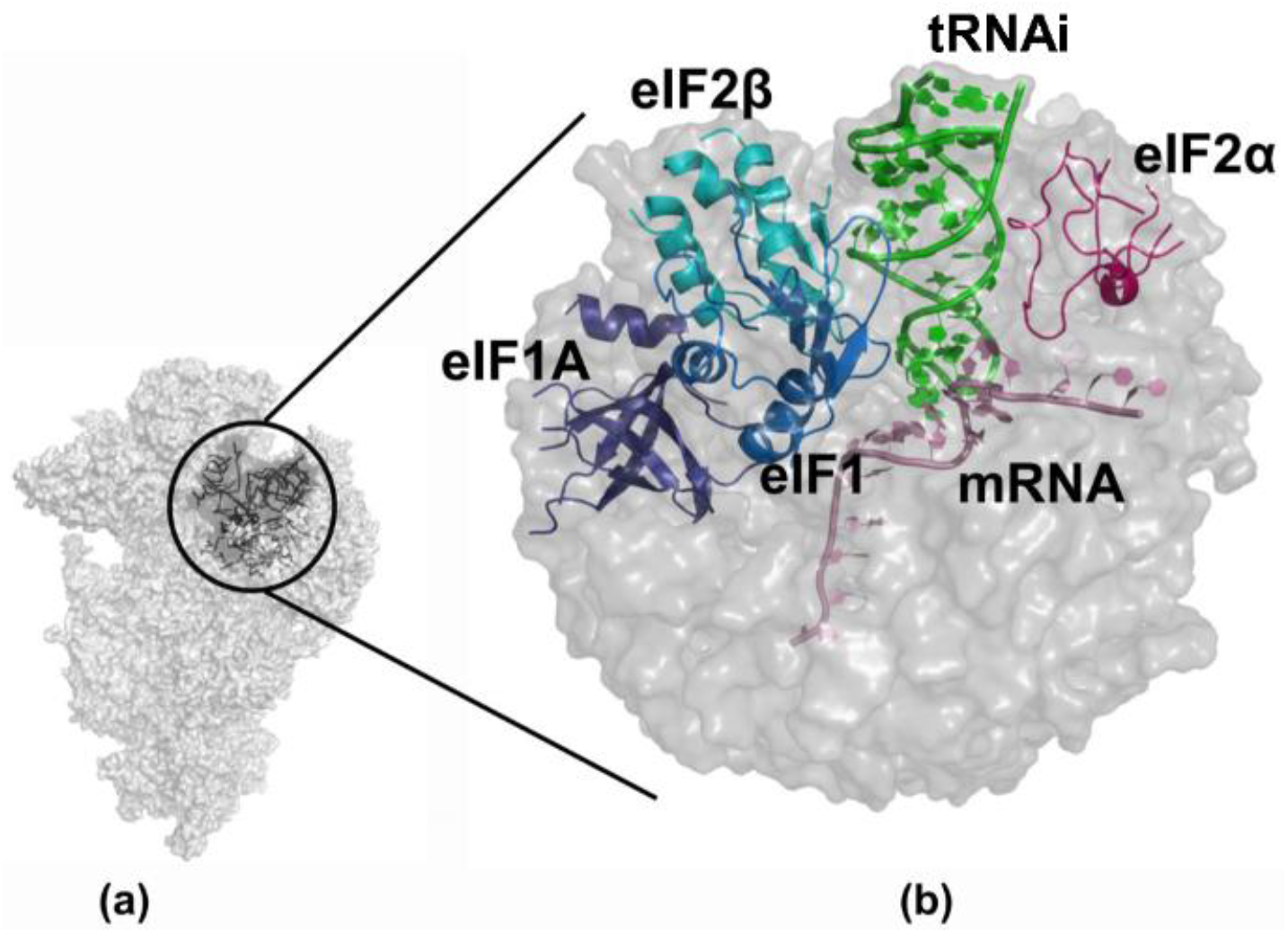
Schematic representation of the simulation sphere. (a) The ribosomal 48S PIC in the open complex is shown in surface representation, and the simulation sphere around P site is encircled (b) In the zoomed view of the simulation sphere, the portions of initiation factors eIF1, eIF1A, eIF2α and eIF2β, the initiator tRNA and mRNA are shown in cartoon representation. The AUG codon at the P site of mRNA was used for calculating free energy for cognate start codon: anticodon (AUG: UAC) interactions. The codon in P site was mutated to different codons to calculate free energy for respective mutant codon: anticodon interactions and the relative binding energies with respect to AUG codon were obtained 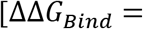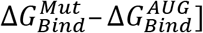

**Fig. 2:**
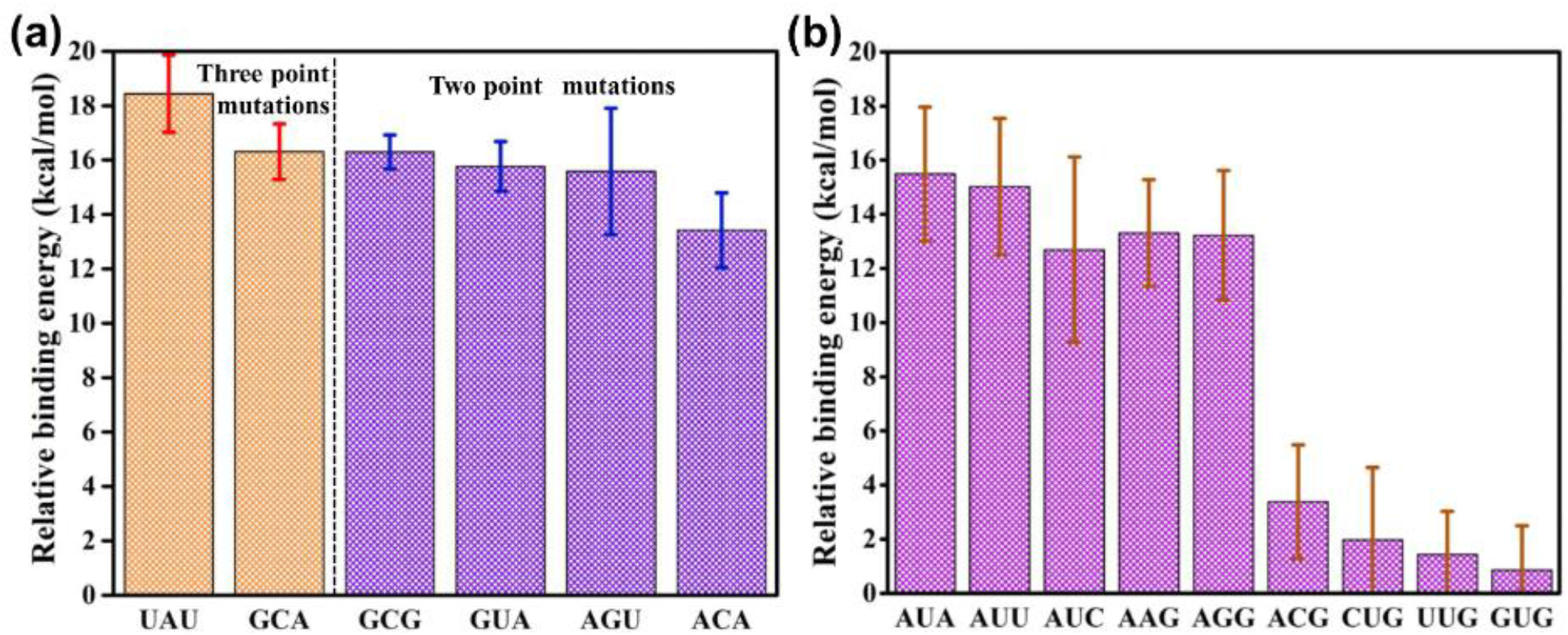
Relative binding energy profiles of (*a*) 3- and 2-point, and (*b*) 1-point mutations of the start codon. 3- and 2-point mutations, where all three or any two nucleotide bases of the start codon are mutated, show a high energy penalty. In 1-point mutation, where any one base of the start codon is mutated, five mutations show a higher energy penalty, whereas four mutations show a low energy penalty. The error bars indicate the standard error obtained from the mean of four independent simulation runs.

#### 3.1.1. Non-cognate codons (two or three mismatches with the tRNAi anticodon)

Out of twenty-seven possible triplets with mutations from AUG at all three positions (3-point mutations), relative binding energies were calculated for two such triplets selected at random, ensuring that both transition (AUG →GCA) and transversion (AUG →UAU) mutations at all three positions were represented. The calculated free energy for cognate start codon: anticodon (AUG: UAC) interactions was used to derive the net relative binding energies, which were obtained by subtracting the respective binding energies of the non-cognate states, respectively 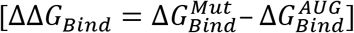. The calculated relative energies for the two non-cognate codons range from 16 to 18 kcal/mol (Fig. 2a). In the average MD simulation PDBs with GCA and UAU, no base pairing was observed between the codon and the anticodon (Supplementary Fig. S2C). We consider it very likely that other non-cognate codons with 3-point mutations have similar high relative binding energies because no base pairing between codon: anticodon is expected in these non-cognate codons as well. Such very high energetic penalties indicate the ability of the 48S PIC to preferentially select AUG over 3-point mutations (GCA and UAU) in its open state.

Next, we carried out similar analyses on four triplets (ACA, GUA, GCG and AGU) with mutations from AUG at two positions, which were selected from all twenty-seven possible 2-point mutations to include transition mutations in codon positions 1 and 2 (GCG) or 1 and 3 (GUA), and both transition and transversion mutations at positions 2 and 3 (ACA and AGU). The relative energies of the 2-point mutations also have high energetic penalties in the range of 13 to 16 kcal/mol (Fig. 2a), further indicating the ability of the 48S PIC to discriminate against non-cognate codons in its open state. Again, in the average MD simulation PDBs with 2-point mutations, no base pairing was observed between the codon and the anticodon (Supplementary Fig. S2B).

The results suggest that the 48S PIC in its open conformation can reject codons with two or three mismatches with the tRNAi anticodon, thereby discriminating most non-AUG codons encountered in the P site during scanning. This would obviate the requirement for conformational switching to the closed state in order to reject the majority of all non-AUG triplets, which might help explain how accuracy is maintained at a high speed of scanning of 5’ UTRs by 48S PIC.

#### 3.1.2. Near cognate codons (one mismatch with the tRNAi anticodon)

We next calculated the free energy perturbations from simulations examining all nine possible near-cognate triplets (GUG, UUG, CUG, AGG, AAG, ACG, AUA, AUU and AUC) with a mutation from AUG at one position (1-point mutation). Intriguingly, the calculated relative binding energies fall into two distinct groups. The first group, consisting of triplets AUA, AUC, AUU, AAG and AGG, showed higher energy penalties in the range of 12-16 kcal/mol (Fig. 2b). The remaining triplets ACG, CUG, GUG, and UUG, forming the second group, showed only moderate penalties of 1-3 kcal/mol (Fig. 2b). Base pairing between codon: anticodon is observed in the latter group having low penalties, whereas it was absent in the former group with high energy penalties (Supplementary Fig. S2A). These results agree with *in vitro* and *in vivo* studies wherein codons in the second group support the highest non-AUG initiation [30,31,32,33,34,35,36,37,38].

The results indicate a strong preference for ‘G’ in the third position of the triplet to achieve the lowest energy penalties, as any of the single-point mutations involving this base (in triplets AUA, AUU, or AUC) conferred much higher energy penalties (Fig. 2b). Moreover, the single-point transversion mutations at the second position (U) to either purine (i.e., AUG→AAG and AUG→AGG) are not tolerated. Likewise, they confer large relative binding energies, which was not observed for the transition mutation of U→C in the AUC triplet (Fig. 2b). Interestingly, both transversion and transition mutations at the first base seem to be tolerated (in triplets CUG, UUG, and GUG), which implies the inability of the PIC in its open conformation to recognize the first nucleotide of the AUG start codon efficiently.

In brief, the calculations suggest that the scanning PIC can reject all but 4 of the 63 non-AUG codons in its open conformation. For the four near-cognate codons with much lower energy penalties, ACG, CUG, GUG, and UUG, the 48S PIC may then proceed to the closed conformation and execute a second accuracy check in order to reject these near-cognate triplets and achieve stringent AUG selection.

#### 3.1.3. Role of eIFs in scanning

To gain insight into the roles of initiation factors in start codon recognition in the open conformation of the 48S PIC, we carried out three sets of simulations in the presence and absence of particular eIFs, including eIF1, the combination eIF1 and eIF1A, eIF2α and eIF2β. In each set of simulation mRNA with cognate (AUG), 1^st^-position near-cognate (GUG), or 3^rd^-position near-cognate (AUA) start codon was used. It may be noted that GUG shows a low energy penalty whereas AUA falls in a high energy penalty group (Fig. 2b)

In the absence of eIF1, the relative binding energy for AUA was found to be 8.4 kcal/mol (Fig. 3), much less than that in its presence (15.5 kcal/mol). For GUG, in contrast, relative binding energy of only ~1 kcal/mol over AUG was observed in the presence of eIF1, and a similar value of 2 kcal/mol was observed in the absence of eIF1 (Fig. 3). This suggests that eIF1 has a role in the discrimination against the 3^rd^-position near-cognate AUA triplet in the open PIC, such that the energy penalty for AUA versus AUG is reduced in the absence of eIF1. By contrast, eIF1 does not seem to discriminate against the 1^st^-position near-cognate GUG triplet in the open PIC, which might help to explain the minimal energy penalty for GUG in the scanning PIC with eIF1 present.

**Fig. 3:**
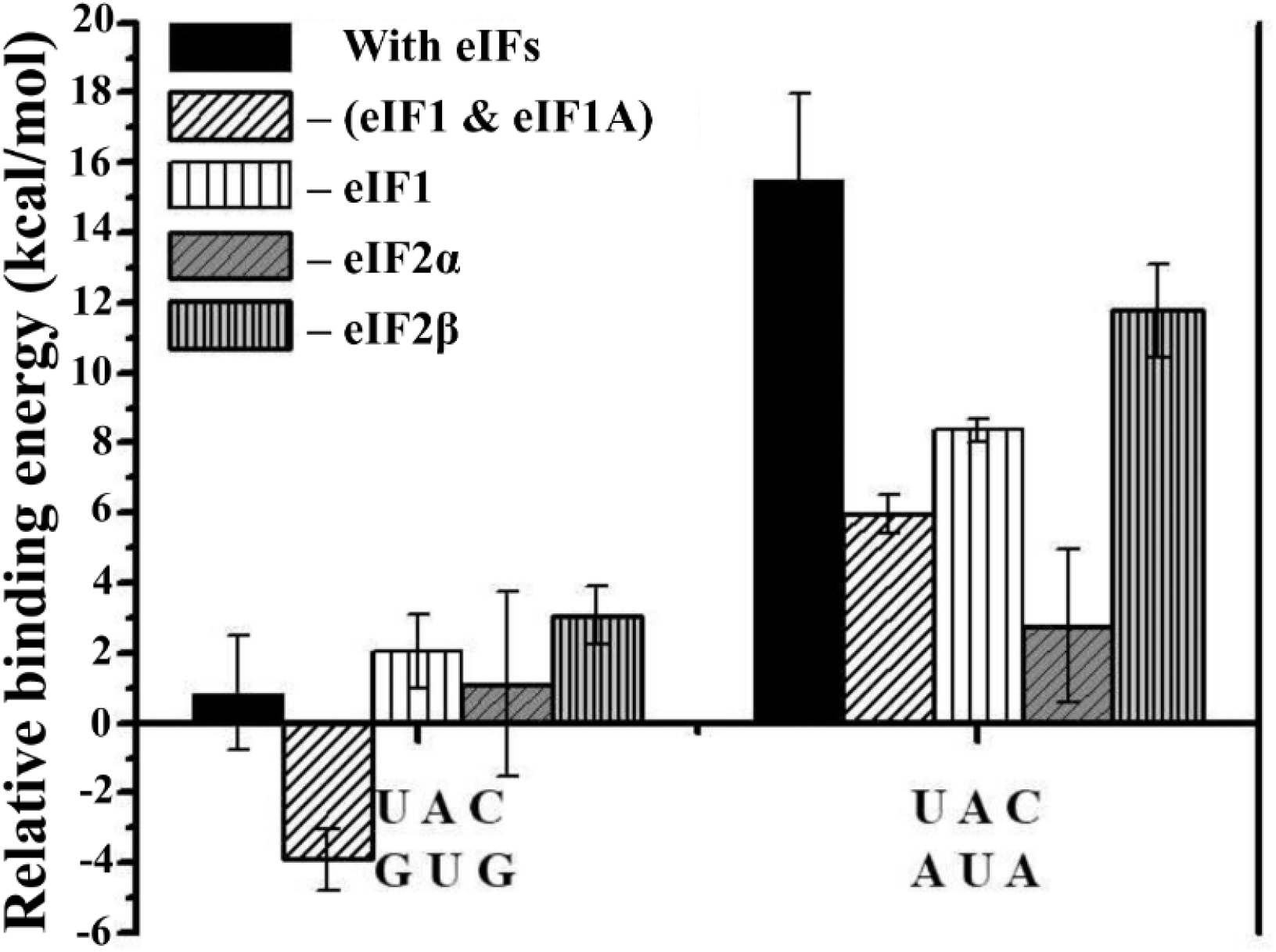
Relative binding energy profile of codon-anticodon interactions from GUG and AUA with respect to AUG simulation runs in the presence and absence of different eIFs. AUA shows much lower relative binding energy in the absence of eIF1 alone, eIF1 and eIF1A or eIF2α. Error bars indicate the standard error obtained from the mean of four independent simulation runs.

eIF1 is known to increase the fidelity of start codon selection by opposing both transition to the closed state and subsequent Pi release at non-AUG codons [3,39,40]. This is achieved through its interaction with eIF2β exclusively in the open complex [19] and by imposing a steric block that prevents the codon: anticodon duplex from achieving the PIN conformation [15,19]. The latter provokes displacement of eIF1 from its original position at the P site on AUG recognition in the closed complex as a prelude to its subsequent dissociation from the PIC [15,19,41]. As a result, eIF1 mutations that decrease its abundance, weaken its interaction with eIF2β or the 40S subunit or diminish its clash with tRNA_i_, all allow inappropriate rearrangement to the closed state at near-cognate UUG codons in vivo [19,42,43,44]. In the open conformation of the 48S PIC, the mRNA channel is widened, and eIF1 is observed in its original position at the P site, exerting no steric hindrance to the codon: anticodon duplex [19]; however, as discussed later, β-hairpin loop-1 of eIF1 in the P site interacts with the codon in average MD structure.

Simulations conducted in the absence of both eIF1 and eIF1A gave results similar to those observed in the absence of eIF1 alone for the AUA triplet, as the relative binding energy was reduced from ~15 kcal/mol to 6 kcal/mol, which is only slightly lower than that in the absence of eIF1 alone (~8.4 kcal/mol). These findings suggest that eIF1A also exerts some discrimination against AUA in the scanning PIC independently of eIF1. Surprisingly, GUG showed a much lower relative binding energy (−3.9 kcal/mol) in the absence of both factors, indicating that it may be preferred over AUG in such a scenario.

Intriguingly, deleting eIF2ɑ from the system conferred a marked reduction in relative free energy in the case of AUA from ~16 kcal/mol to 2.8 kcal/mol, while that of GUG remained virtually unchanged at 1.1 kcal/mol (Fig. 3). eIF2ɑ is bound at the at E-site, and it interacts with both tRNA_i_ and mRNA upstream of the start codon in the P_IN_ state [15]; however, its role in the fidelity of start codon selection is not well understood. The absence of eIF2ɑ from the system, as discussed later, gives more dynamic flexibility to mRNA for base-pairing with the tRNAi anticodon in the P site (Supplementary Fig. S3), which might account for the lower energy penalty for the AUA codon.

Eliminating eIF2β results in a small decrease in energy penalty for AUA (from ~16 to 11.8 kcal/mol), but an increase for GUG (from 0.86 to ~3 kcal/mol). As described above for eIF1, these perturbations might be expected to increase initiation at AUA codons but decrease it at GUG or UUG codons. As noted above, mutations expected to weaken eIF2β interaction with eIF1 in the open conformation of the 48S PIC increase UUG initiation [19], which can be explained by increased rearrangement to the closed complex at UUG codons that presumably outweighs the slightly increased discrimination at UUG codons in the open complex predicted from our results in Fig. 3. Considering that eIF2β is also in the vicinity of eIF1A and the tRNAi anticodon stem-loop in the py48S-open complex [19], the changes in relative binding energy for AUA and GUG in the absence of eIF2β (Fig. 3) might indicate an indirect role in codon selection in the open state by virtue of its interaction with eIF1, eIF1A, or tRNAi.

### 3.2. Structural insights on codon selection in the scanning conformation of the 48S PIC

The calculated relative binding energies indicate that the open 48S PIC complex can discriminate against many non-cognate as well as near-cognate codons. Interestingly, only four codons, all near-cognates, show a low penalty, and the contributions of eIFs to the larger penalty incurred with the AUA triplet was revealed by the reductions in this penalty observed in their absence. Average MD structures taken from the individual runs, as well as the extracted PDBs from the simulation runs, were analyzed to figure out the mechanism of codon selection by the 48S PIC in its open conformation.

#### 3.2.1. Recognition of AUG in the open state

The available cryo-EM structures of yeast 48S PIC in an open conformation have an AUC start codon in the mRNA where only the A and U bases of the codon in the P site could be clearly observed [19,20]. Thus, in the absence of a structure of the 48S PIC in scanning conformation with an AUG start codon, which has not yet been captured experimentally for structural determination, the average MD structure with an AUG codon provides insights into recognition of the correct start codon in the open state (Figs. 4a and 4b). This structure reveals a stable codon: anticodon interaction at the P site (Fig. 4a). The position of the mRNA in the channel as well as the codon at the P site in the average MD structure in the open state is different from that observed in the closed conformation [21] (Supplementary Fig. S4a). Hence, the mRNA including the codon at the P site, is repositioned as the mRNA channel is narrowed during the transition from open to the closed state. Interestingly, similar observation in the change of mRNA and start codon position from open to closed conformation of PIC was also observed in bacterial translation initiation [45].

**Fig. 4:**
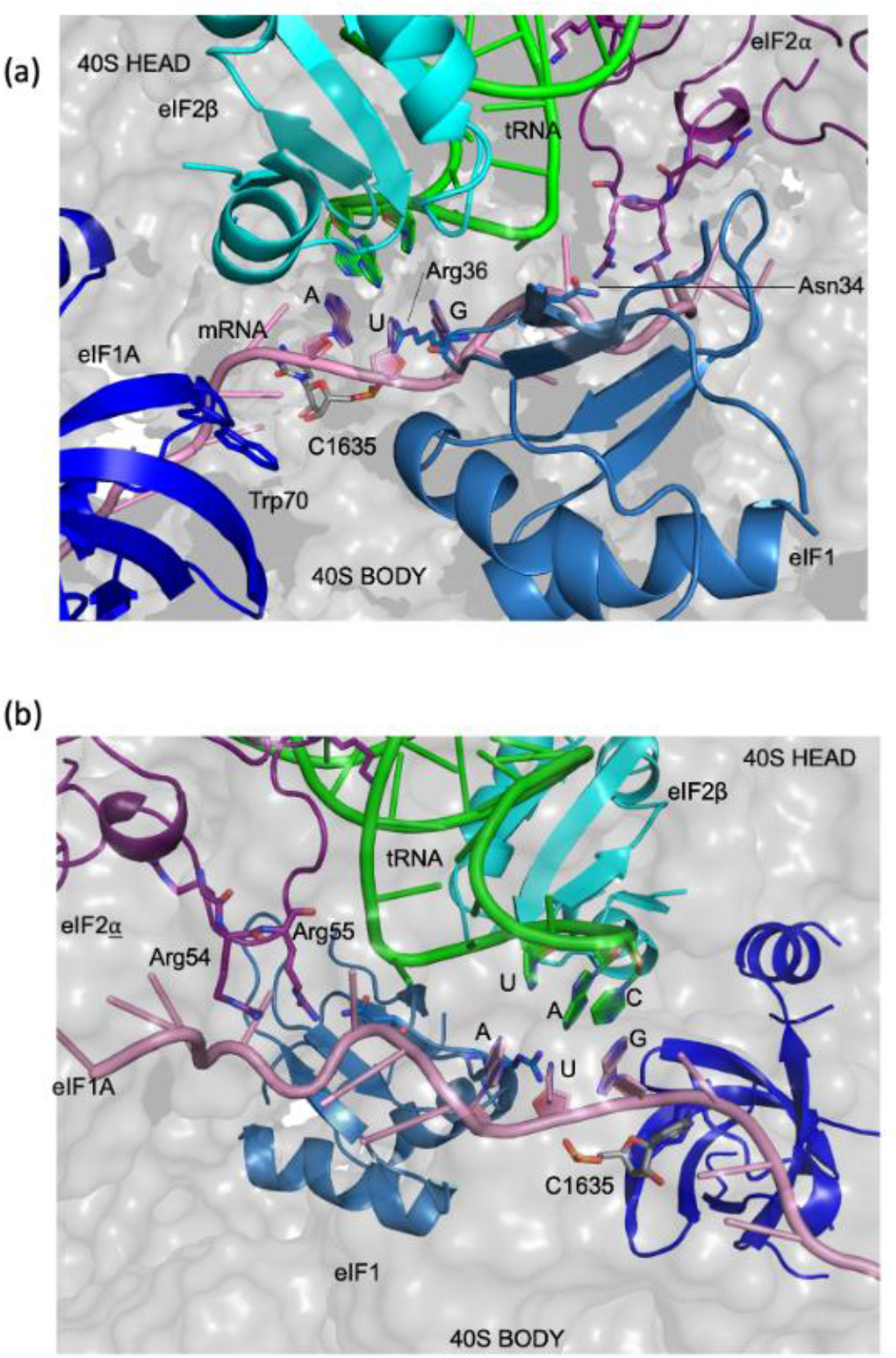
Recognition of the start codon AUG by 48S PIC in the open conformation. (*a* & *b*) The key players involved in the codon-anticodon interactions are shown in cartoon representation and the corresponding residues in stick representation in average MD structure. Arg36 of eIF1 (pale blue) interacts with the first two codon bases, while Trp70 of eIF1A (blue) points towards the mRNA, providing stacking interactions. eIF2α, shown in purple colour, contributes a series of Arg residues (*b*), which interact with mRNA. eIF2β (cyan) interacts with both tRNAi and eIF1 at the P site. The tRNAi and mRNA are shown in green and pink, respectively.

In the average MD structure with AUG codon, Arg36 present in β-hairpin loop-1 of eIF1 interacts with the first two nucleotides (A and U) of the codon (Fig. 4a), whereas this loop of eIF1 has no interaction with the codon bases in the py48S-open and py48S-open-eIF3 structures [19,20]. Notably, the main chain amino group of Arg36 interacts with ‘U’ of CAU anticodon of the codon: anticodon duplex in py48S and py48S-closed complexes [15,19,20]. Moreover, Asn34, also present in β-hairpin loop-1 of eIF1, interacts with the first nucleotide A of the codon in another average MD structure with AUG codon (Supplementary Fig. S4b); however, in this case, the interaction with Arg36 is not observed. Asn34 interaction with anticodon was seen only in the closed state of 48S PIC [15,19,20]. Thus, interactions observed with Asn34 and Arg36 in the average MD structures seem to indicate how these residues hold onto the codon: anticodon duplex in the open state with wider P site, and these residues continue to interact with codon: anticodon duplex even after the transition to the closed state.

No direct contacts of eIF1A with the mRNA were found in the average MD structure with an AUG codon. Trp70 of eIF1A was observed to stack with +4 nucleotide of the mRNA in the closed state of py48S and py48S-closed complexes [14,19,20]. However, in the open state of 48S, no interaction between Trp70 and mRNA is observed in the simulation run with AUG (Fig. 4). It is likely that as the mRNA channel is narrowed during the transition from the open state to the closed state, the base of the +4 nucleotide of the mRNA flips out to stack with the Trp70 of eIF1A. The base of rRNA residue C1635 provides stacking interactions to the third codon base (Fig. 4), which in turn would stabilize the base-pairing between codon: anticodon. C1635 was also observed stacking the third codon base pairing with the anticodon in the 48S PIC structures with AUG codon in the closed state [15,19,20].

Furthermore, a series of Arg residues in a loop of eIF2ɑ and various residues of ribosomal proteins uS7, uS11 and eS28 could be observed interacting with mRNA upstream of the AUG codon in the P site (Fig. 4 and data not shown). These are similar to the interactions observed in the P_IN_ state of py48S-closed, but were not observed in the cryo-EM map of py48S-open, probably due to lack of distinct mRNA density in the widened mRNA channel. Arg54 and Arg55 of eIF2ɑ form stable hydrogen bonds with the− 3 and− 4 bases of mRNA (Fig. 4), which belong to the Kozak sequence. Thus, it appears that eIF2ɑ interacts with mRNA and binds to the Kozak sequence upstream of AUG in the P site even in the open conformation of the 48S PIC. eIF2β is sandwiched between tRNA_i_ and eIF1 as observed in py48S-open-eIF3 complex, and forms hydrogen bonds with both tRNA_i_ and eIF1. Thus, the average MD structure with an AUG codon provides insights into the interaction of the mRNA, including the start codon with eIFs and ribosomal proteins and rRNA in the open state.

#### 3.2.2. Discrimination of non-AUG codons in the open state

The systems with low energy penalty have a stable tRNA_i_ during the course of the simulation, which plays a prominent role in forming and maintaining the codon-anticodon interactions. Whereas in scenarios of high energy penalty, the tRNA_i_ does not stabilize during the course of the simulation, and the anticodon stem-loop (ASL) was observed to be more dynamic (Fig. 5), thus breaking the codon-anticodon interaction. The β-hairpin loop-1 of eIF1 was found to be dynamic as well. However, apart from loop-1, the overall structure of ‘eIF1’ was stable and was superimposable onto one another with a relatively low root mean square deviation (RMSD) score of ~0.8 Å in multiple simulation runs.

**Fig. 5:**
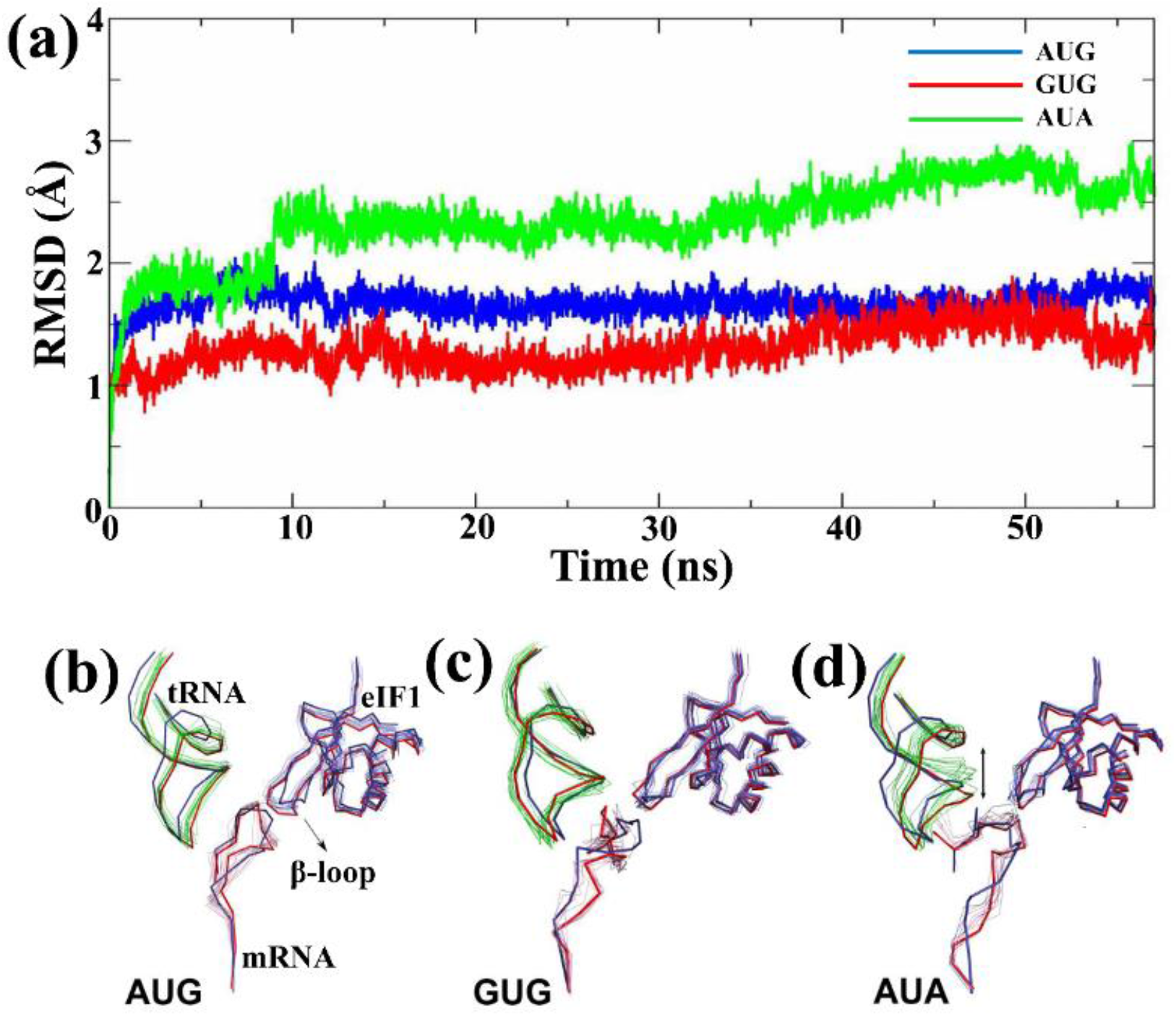
RMSD of tRNAi backbone and average PDBs. (*a*) RMSD of tRNA_i_ backbone from three independent simulation runs of AUG (black), AUA (green) and GUG (red), respectively. (*b-d*) Snapshots of the average PDBs extracted from the last 5 ns of the simulation trajectories from the respective (AUG, GUG and AUA) simulation runs are shown in ribbon representation. The PDBs were superposed based on rRNA stretch from the 40S body. The anticodon stem-loop of tRNA_i_ (green) is more stable for (*b*) cognate and (*c*) near-cognate codons, whereas it is more dynamic for (*d*) non-cognate codon. The starting and final structures for each individual runs are shown in pink and blue, respectively.

The loop-1 makes interactions with the first two nucleotides in the codons in multiple runs with different codons at the P site (data not shown). Similarly, the interaction of loop-1 with A and U of the anticodon is also observed in multiple runs in the open state (data not shown). However, no interaction with the nucleotide at the third position of codon or with corresponding C of anticodon was observed for eIF1. Thus, how eIF1 influences the selection at the third position of the codon and, at the same time, tolerates mismatch at the first position in the open state is not clear. We suggest that the loop-1 of eIF1 by interacting with codon: anticodon in the open state reduces the available space at the P site for any relaxed association between mRNA and tRNA even in widened mRNA channel, thereby discriminating against most codons. Codons with a mismatch in 1^st^ position (GUG, CUG and UUG) and codon with a pyrimidine in 2^nd^ position, i.e., ACG, are tolerated even in the presence of eIF1. Further, the third position interaction is stabilized by stacking with C1635 (discussed below), and the presence of loop-1 of eIF1 and rRNA C1635 seems to ensure the selection of correct base-pair at the third position. In the absence of eIF1, the available space at P site is increased, thereby decreasing the energy penalty even for third position mismatch (Fig. 3).

In simulations of the open PIC complex containing an AUG start codon, Arg54 and Arg55 of eIF2α stably interact with the (−3/-4) bases upstream of the AUG codon in the P site (Figs. 4b and 6a). In the AUA simulation, by contrast, Arg54 and Arg55 are dynamic, shifting its position to interact alternatively with tRNA_i_ or mRNA during the course of the simulation (Fig. 6c and Supplementary Fig. S5). Thus, in this scenario where the codon: anticodon interaction is not stable, these residues can frequently lose their interaction with the mRNA, which might allow the mRNA to move and bring the next triplet of nucleotides into the P site. Interestingly, in GUG simulations, the dynamics of these Arg residues of eIF2α appears to be intermediate between the AUG and AUA (Fig. 6b and Supplementary Fig. S5). It seems that Arg54 and Arg55 of eIF2α stabilize the mRNA in place in the widened channel of the open conformation of 48S PIC in the scenario of codon-anticodon interaction at the P site. In the absence of eIF2α from the system, the mRNA is found to be more flexible, thereby allowing near-cognate AUA to base-pair with the tRNAi anticodon in the P site (Supplementary Fig. S3).

**Fig. 6:**
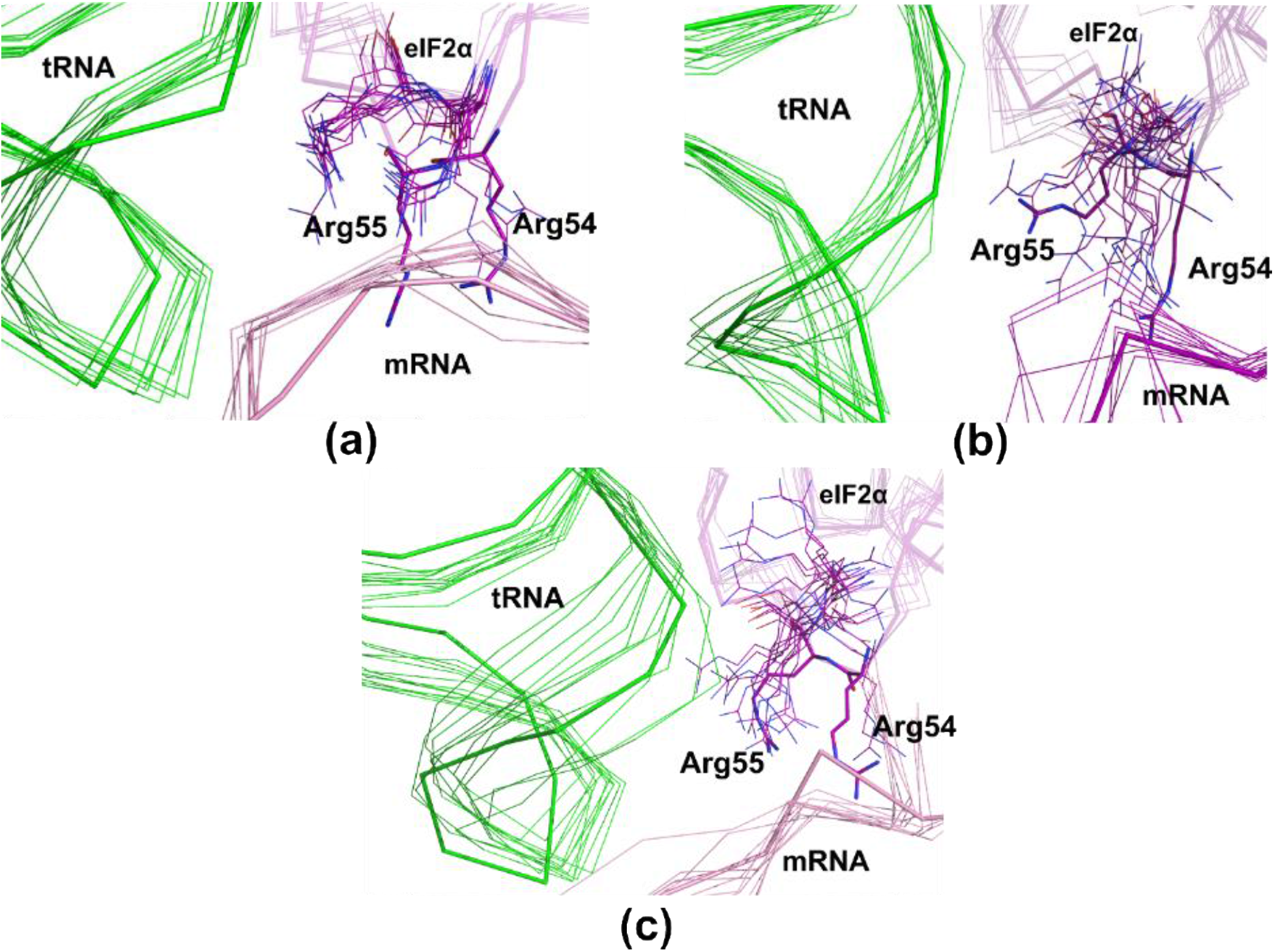
Dynamics of Arg54 and Arg55 of eIF2α from the (*a*) AUG, (*b*) GUG and (*c*) AUA simulation runs. The frames were extracted from the respective trajectories at 5 ns intervals. The starting structures for the runs are shown in thicker ribbon representation. The dynamics of mRNA and Arg residues were more prominent for AUA.

eIF2β is positioned between tRNAi and eIF1 and forms hydrogen bonds with both tRNA_i_ and eIF1 (Fig. 4). As mentioned above, the eIF2β-eIF1 interaction, found exclusively in py48S-open, appears to increase accuracy by maintaining the scanning conformation of the PIC at near-cognate UUG codons [19]. Interestingly, comparing simulations conducted with either AUG, GUG, or AUA codons, we observed a moderate decrease in the number of hydrogen bond interactions at the eIF2β-eIF1 interface for GUG versus AUA, and a larger reduction for AUG versus both GUG and AUA (Supplementary Fig. S6). This observation may indicate that the system is shifting towards the closed state upon correct start codon recognition.

The rRNA residue C1635 stacks the third codon base, thereby stabilizing the base-pairing between codon-anticodon in the case of AUG simulations, as discussed above. In GUG simulations, C1635 is also observed to stack the third codon position (Supplementary Fig. S2). Whereas in the case of near-cognate AUA and non-cognate codons AGU, ACA, and UAU where no base-pairing is observed between codon-anticodon, C1635 no longer stacks the third base of the codon (Supplementary Figs. S2b). The stacking of C1635 with codon-anticodon in also observed in py48S and py48S-closed complexes [15,19]. Thus, this interaction in AUG and the near cognate GUG codon simulations may indicate that the 48S PIC is preparing to transit towards the closed conformation upon codon-anticodon base-pairing in the open state.

## 4. Conclusions

Computational simulation studies have been used to bring out the detailed energetics and intermolecular interactions involved in various steps in protein synthesis [46,47,48,49,50,51,52]. Recently, MD simulations studies were done to study codon selection in translation initiation in the closed conformation of the 48S PIC [16,17]. We have carried out extensive MD simulations studies to compare the relative binding energy of the tRNA_i_ with different non-cognate codons with respect to start codon, i.e., AUG in the open conformation of 48S PIC. A simulation sphere of 40 Å radius centered on ‘A’ nucleotide of the anticodon CAU of tRNA_i_ from the py48S-open-eIF3 complex was generated and used for the simulation studies to provide crucial insights into the energetics of start codon selection by the 48S complex in its scanning conformation. The cryo-EM structure of the py48S-open-eIF3 complex (PDB ID: 6GSM) [20] does not have densities corresponding to the N- and C-terminal tails of eIF1A and hence these are not accounted for in this study. While the NTT plays a role in stabilizing the closed conformation of 48S PIC, the C-terminal tail (CTT) of eIF1A extends into the P site [53] and stabilizes the open conformation of the 48S PIC [54]. Recognition of AUG would lead to the removal of eIF1A-CTT from the P site [55]. In the absence of eIF1A-CTT in the structure of the py48S-open-eIF3 complex [20], this study is also carried out without taking into account eIF1A-CTT. Hence, how eIF1A-CTT stabilizes POUT conformation and mutations in CTT facilitate POUT to PIN transitions at near cognate codon [54] remains to be figured out.

Our extensive MD simulation studies with cognate, near and non-cognate codons in 48S PIC gave crucial insights into the energetics of start codon selection by the PIC in its open state. The result indicates the ability of tRNA_i_ to preferentially base pair with the cognate start codon (AUG) in POUT or open state. Base-pairing of codon: anticodon stabilizes the tRNA_i_ in widened mRNA channel and β-hairpin loop-1 of eIF1 monitors the codon: anticodon interaction. eIF2α interacts with mRNA at E site, preparing the 48S PIC in the open state to change its conformation to the closed PIN conformation.

The tRNA dynamics in the widened P site in the open state seem to drive the selection of the codons. Stable codon: anticodon interaction, as in the case of AUG and GUG codons, lead to a stable tRNA during the simulation. The β-hairpin loop-1 of eIF1 protrudes into the P site and interacts with the AUG codon. Though it poses no steric hindrance to the codon: anticodon in the widened P site in the open state, it reduces the available space for an incorrect association between codon and anticodon. However, in the closed state, eIF1 poses a steric hindrance to the complete accommodation of the codon: anticodon interaction in the narrow P site and exerts stricter criteria for recognition of cognate codon: anticodon interaction. This study suggests that eIF1 plays a role in codon selection in the open conformation of 48S PIC as the simulation runs in the absence of eIF1 leads to a decrease in the energy penalty for non-AUG codons.

Moreover, this study also suggests that most codons may be discriminated against by the open form of 48S PIC and only four near-cognate codons (GUG, CUG, UUG and ACG) are likely to pass through to the close form by the measure of energy penalty of binding. Subsets of near-cognate codons are start site for initiation and particularly GUG, CUG, UUG and ACG codons have been reported to initiate translation [1,2,3,31,37,56]. In addition, codons AAG and AGG, which shows a high penalty in our study, are also reported not to initiate translation [33,37,56,57]. While AUA, AUU and AUC codons, which shows high penalty in our work, have been shown to act as a start codon in *Neurospora crassa,* their ability to be recognized as alternate start codon is with much lower efficiency [37,58]. Thus, overall the selection of codons by the 48S open form, as suggested by this study, correlates well with the non-AUG codons reported to initiate translation.

Seemingly, the open state of 48S acts as the first step of start codon selection, where a coarse selection is made. It would allow only 4 of 63 non-AUG codons to the next closed state for further fine selection. This arrangement of a coarse selection in the open conformation would ensure a high speed of scanning long 5’ UTR, avoiding the need to change conformations to the closed state for each triplet encountered at the P site. Shuttling back and forth between the open and closed states of 48S PIC, for every new codon in the P site would require a conformational change in various components of 48S PIC, amounting to making and breaking of several interactions between them. Moreover, dynamic switching back and forth between open and closed states may not account for the high speed of scanning by 48S PIC.

The open conformation of 48S PIC would rule out most non-AUG codons, thereby allowing a much more thorough checkup point of only a few near-cognate codons in the closed state. Recognition of correct start codon and accommodation of codon: anticodon in P site in closed state triggers the repositioning and eventually release of eIF1. The vacant site at P site is now occupied by the N-terminal domain of eIF5, which then rechecks the codon in the P site [21]. Thus, allowing the codon: anticodon interaction to be monitored at multiple checkpoints during initiation ensures a more thorough and robust codon selection mechanism. This study suggests how the 48S PIC strikes a balance between the accuracy of codon selection and high speed during the scanning process by employing the open state as a coarse selectivity checkpoint to reject all but a few of the possible codon: anticodon mismatches.

## Author contributions

T.H. and P.K.M. designed research; I.B. and B.G. performed research; I.B., B.G., T.C., and T.H. analyzed data; and I.B., T.C., P.K.M. and T.H. wrote the manuscript.

## Acknowledgement

The authors thank Alan G. Hinnebusch for insightful comments and suggestions. I.B. thanks SERB-National Post Doctoral Fellowship (N-PDF) and B.G. acknowledges Dr. D.S. Kothari Postdoctoral Fellowship (DSKPDF) for financial supports [201718-BL/16-17/0437]. T.C. was supported by postdoctoral fellowship from the DBT/Wellcome Trust India Alliance grant (IA/I/17/2/503313) awarded to TH. TH acknowledges funding from DBT/Wellcome Trust India Alliance (IA/I/17/2/503313), DST-FIST [SR/FST/LS11-036/2014(C)], UGC-SAP [F.4.13/2018/DRS-III (SAP-II)] and DBT-IISc Partnership Program Phase-II (BT/PR27952-INF/22/212/2018). We thank Sahasrat, SERC, and TUE-CMS, SSCU at IISc, Bangalore, India for the computational facilities.

## Supplementary Information

**Supplementary Fig. 1:**
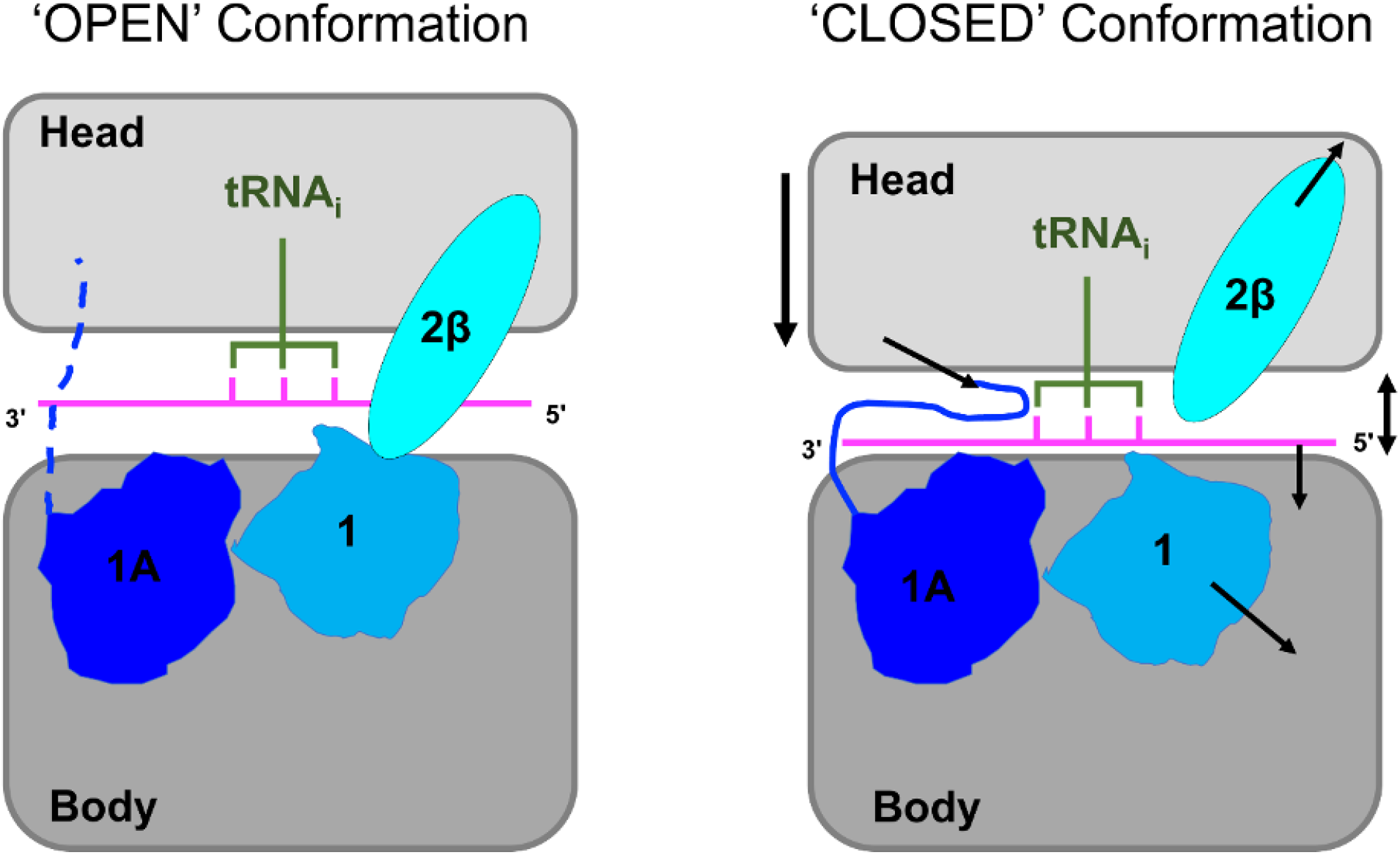
Schematic of open and closed conformation of 48S PICs highlighting the major differences between them. The arrows shown in the closed conformation points out the differences in conformation, namely, 1. the 40S head moves down closer with respect to the body with attendant compression of rRNA helix 28, 2. the mRNA channel and the P site are narrowed, 3. the mRNA and tRNA_i_ positioned in the P site ~7 Å closer to the 40S body compared to that found in py48S-open complex, 4. eIF1 undergoes subtle repositioning away from original binding position on the transition to the closed state to accommodate tRNA_i_ in the PIN conformation, 5. the N-terminal tail (NTT) of eIF1A is observed in proximity to the codon: anticodon duplex, and 6. eIF2β moves away from eIF1 in the closed conformation.

**Supplementary Fig. S2:**
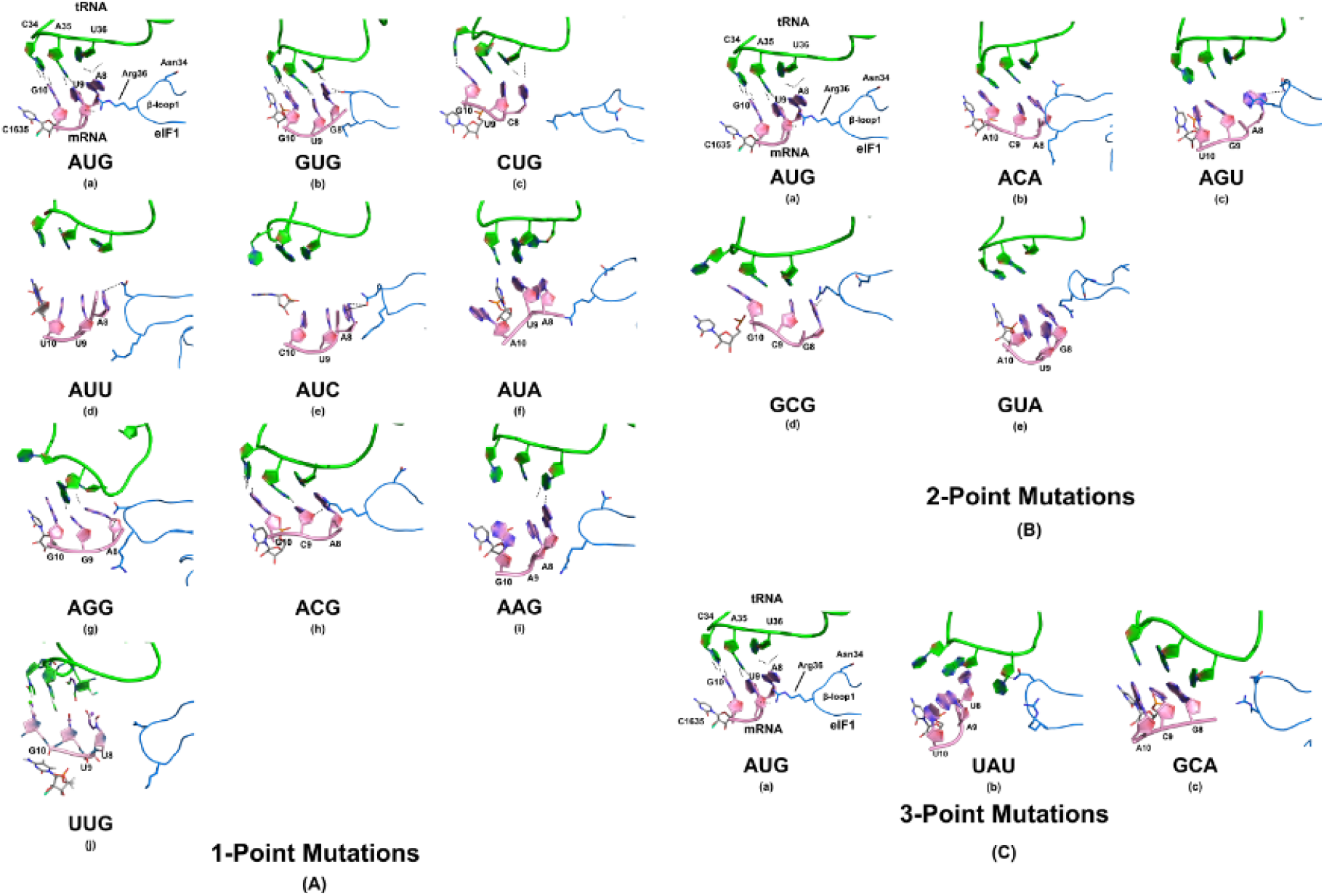
Cartoon representation of codon-anticodon interactions of AUG, (*A*) 1-point mutations, (*B*) 2-point mutations and (*C*) 3-point mutations. tRNA_i_ is shown in green, mRNA in pink, eIF1 in blue, and C1635 of rRNA, which provides the stacking interaction to codon-anticodon in grey colour. Arg36 and Asn34 of eIF1 are shown in stick representations. In AUG, base pairing between codon: anticodon is observed. (A) Among codons with 1-point mutation, in GUG, CUG, UUG and ACG, base-pairing is observed while no base pairing is seen in AUA, AUU, AUC, AGG and AAG codons. No base pairing is observed in no-cognate codons with 3- and 2-point mutations (B and C).

**Supplementary Fig. S3:**
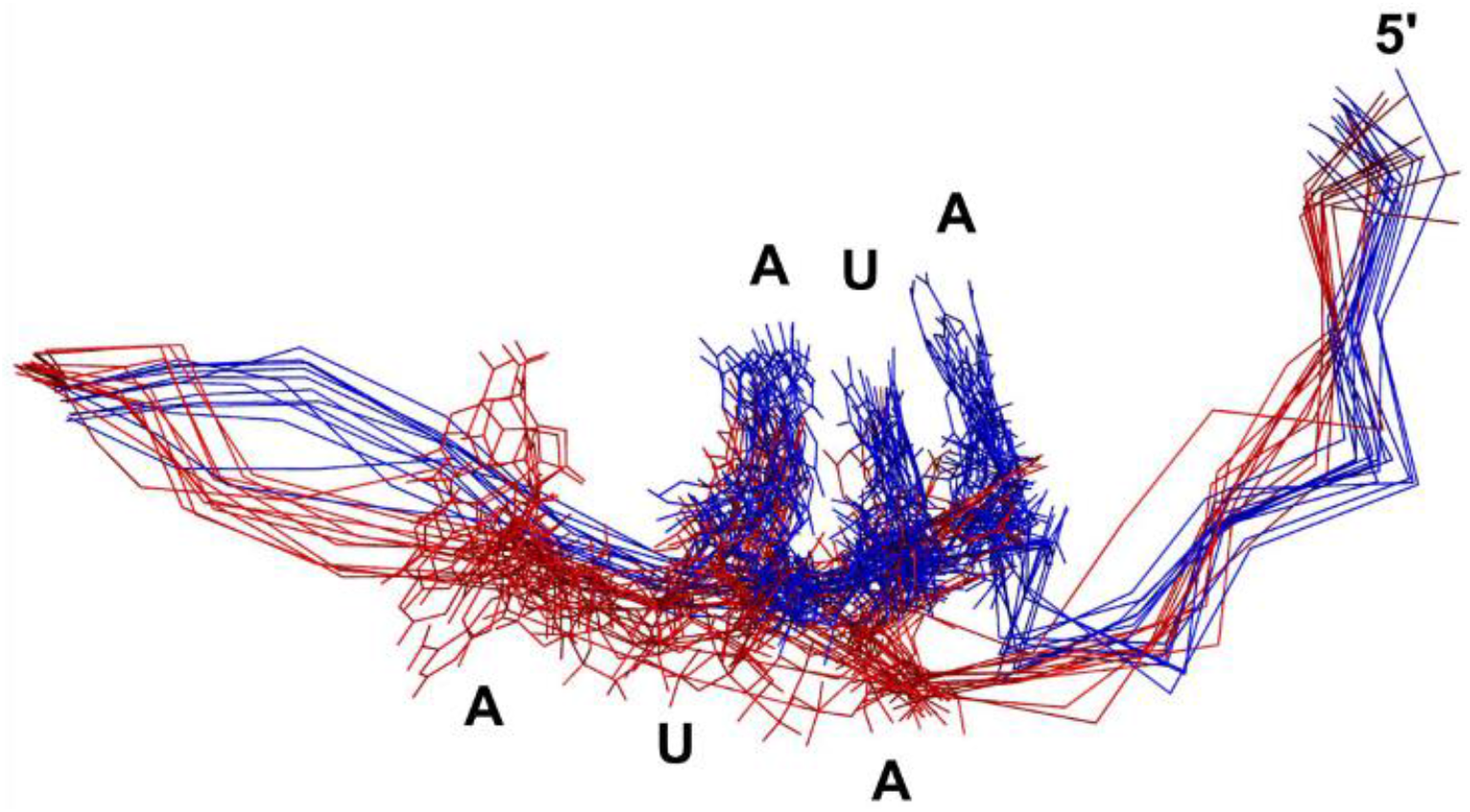
Snapshots of mRNA extracted from the last 5 ns of the simulation trajectories from the AUA simulation runs with (red) and without (blue) eIF2α. The codon AUA is shown to indicate a shift in the codon's position towards the E site without eIF2α (blue) simulation run.

**Supplementary Fig. S4:**
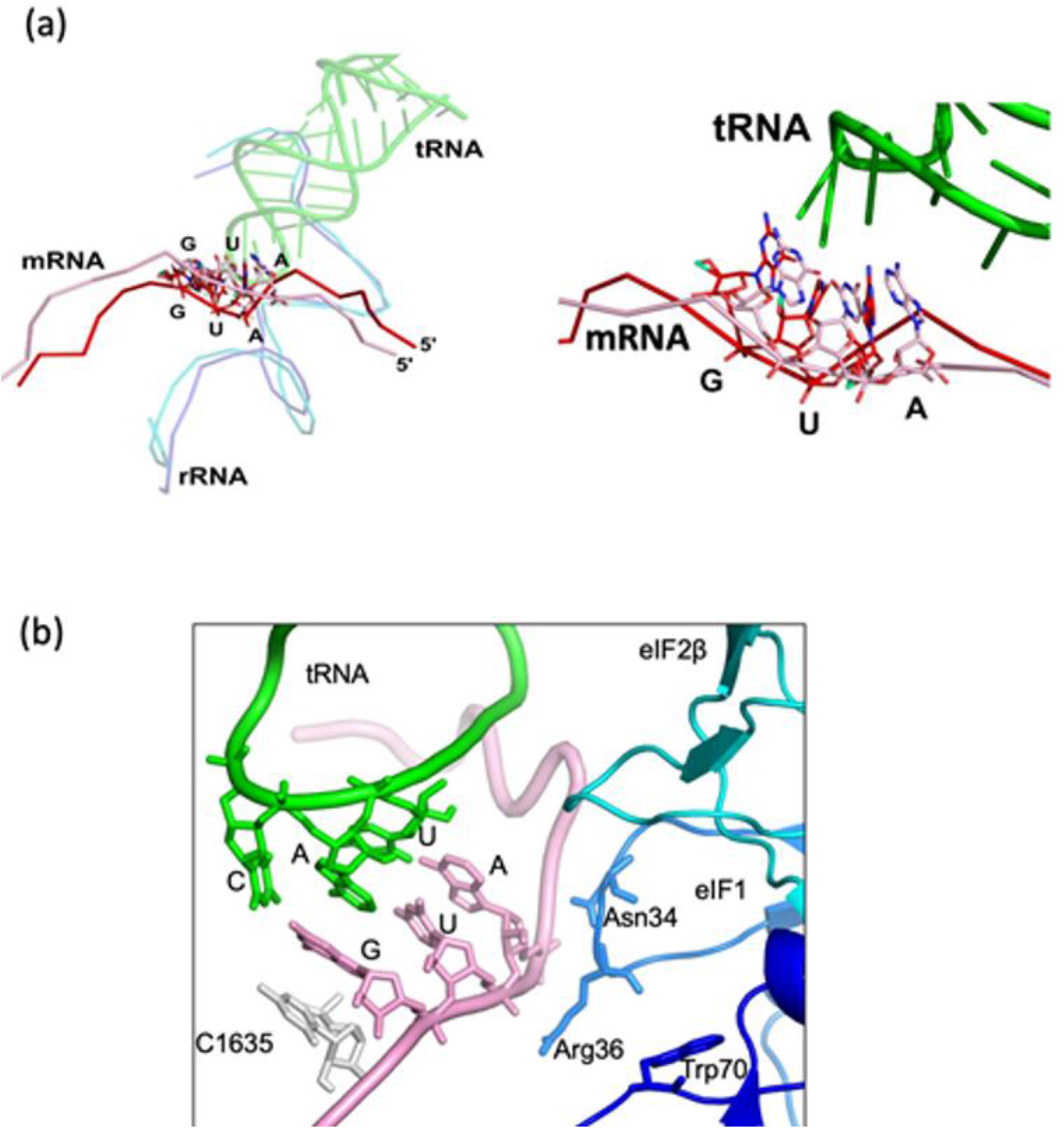
Recognition of AUG start codon during scanning. (a) The mRNA (and start codon) in average MD structure of AUG simulation run in open conformation (red) occupies a different position when compared with mRNA in 48S PIC in closed state (PDB ID: 6FYX (pink). The PDBs were superposed onto each other based on the rRNA (1147-1174; chain 2) from the body of the 40S ribosome. The rRNA from AUG simulation run is shown in dark blue whereas rRNA from PDB ID: 6FYX is in light blue. The tRNA_i_ (green) at the P site is from average MD structure of AUG simulation run in open conformation. An enlarged view of the codon:anticodon is on the left. (b) Asn34 of eIF1 interacts with the first nucleotide A of the codon in another average MD structure with AUG codon.

**Supplementary Fig. S5:**
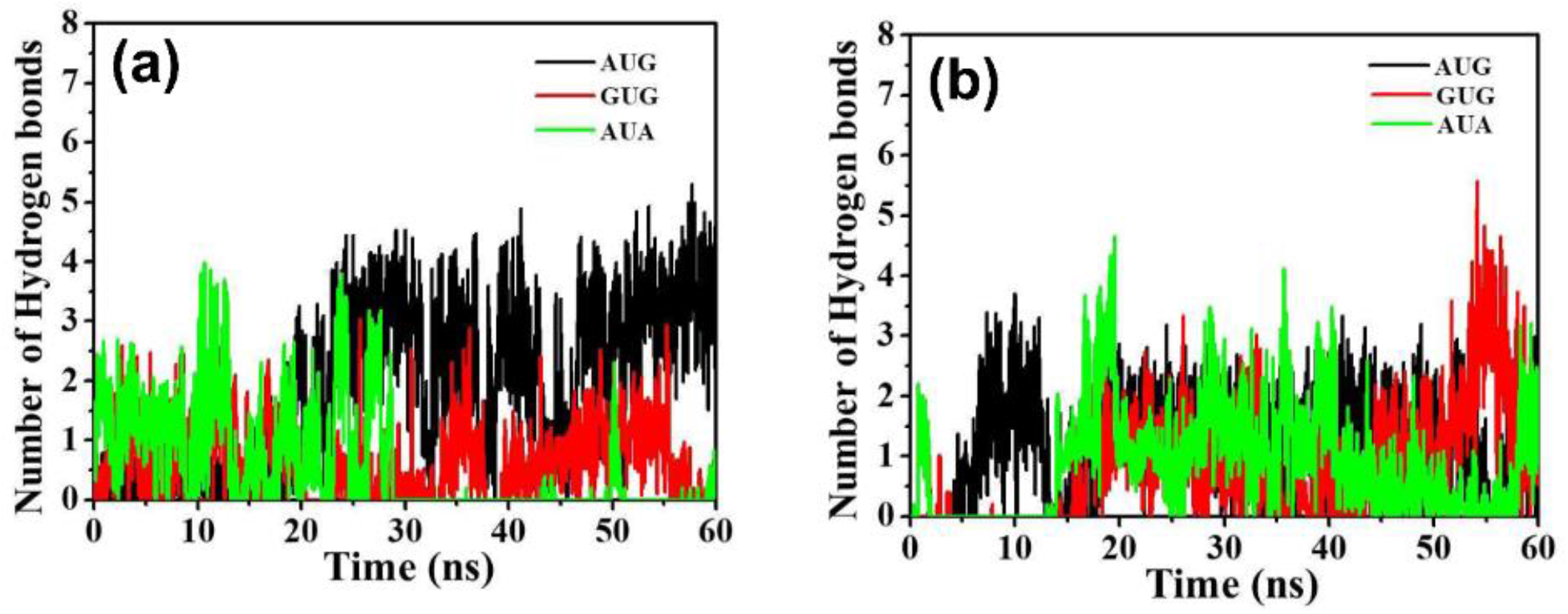
Number of hydrogen bonds between eIF2α residues (*a*) Arg54 and (*b*) Arg55 with mRNA as a function of simulation time for AUG, GUG and AUA runs, respectively. The interactions were stable for AUG and GUG but not for AUA.

**Supplementary Fig. S6:**
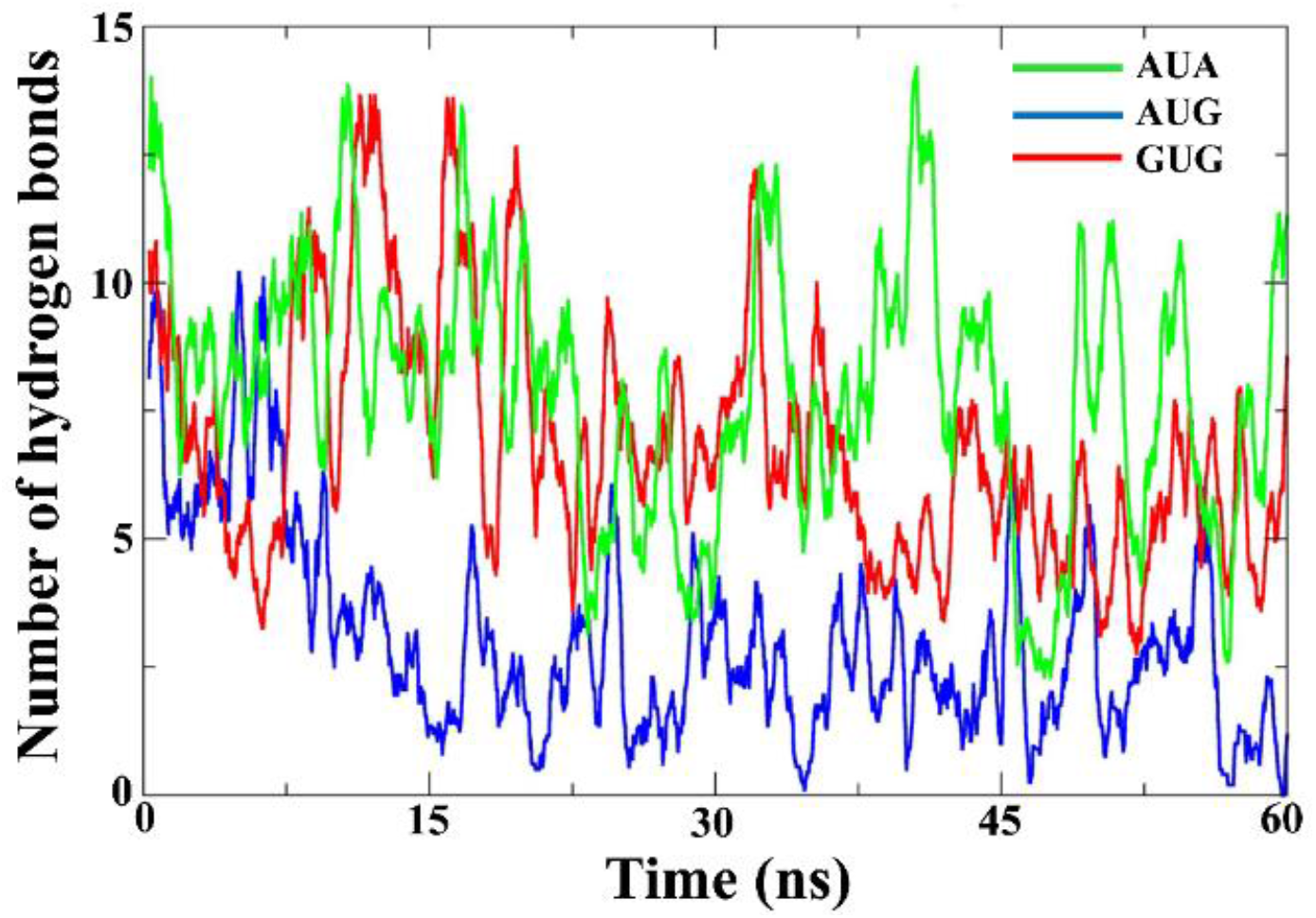
Number of hydrogen bonds between eIF1 and eIF2β interface as a function of simulation time for AUG, GUG and AUA runs. The output from AUA, AUG and GUG simulation runs are show in green, blue and red, respectively. The decrease in the number of hydrogen bonds for cognate codon (AUG) is noticed compared to GUG and AUA codons.

